# Hepatitis C virus genotype homogeneity and lack of major Sofosbuvir resistance-associated substitutions among blood donors in Ethiopia

**DOI:** 10.64898/2026.07.15.738611

**Authors:** Helen Terefe, Tadelo Wondmagegn, Abaysew Ayele, Dawit Hailu Alemayehu, Gashaw Adane, Yimer Demisse, Gizachew Gemechu, Adane Mihret, Tesfaye Gelanew, Andargachew Mulu

**Author notes:** Corresponding author: Helen Terefe, Affiliation: Department of Immunology and Molecular Biology, School of Biomedical and Laboratory Sciences, College of Medicine and Health Sciences, University of Gondar, Gondar, Ethiopia, Email Address, Address: Addis Ababa, Ethiopia /+251922410897.

## Abstract

Hepatitis C virus (HCV) is a hepatotropic virus that causes a spectrum of liver diseases. HCV genetic diversity influences transmission, pathogenesis, prognosis, and response to antiviral therapy. Moreover, resistance associated substitutions (RASs) challenged the existing antiviral treatment. However, data regarding circulating genotypes and RASs are limited in Ethiopia. Therefore, this study aimed to determine HCV genotypes and RASs among apparently healthy blood donors. This study used 97 archived anti-HCV-positive serum samples obtained from apparently healthy blood donors, collected in accordance with national routine blood collection practices. The partial HCV NS5B gene of 45 samples was amplified and sequenced using amplicon-based next-generation sequencing. Genotypes and subgenotypes were determined using NCBI BLAST, Geno2Pheno [hcv], and the Los Alamos HCV database, and a phylogenetic tree as a confirmation. The RASs were determined using Geno2Pheno [hcv]. Of the 97 anti-HCV-positive samples, 45 had detectable HCV RNA, of which 35 yielded high-quality sequences. These strains (n = 35) showed marked homogeneity in NS5B gene sequences: 34 were HCV genotype 4, all subgenotype 4d, and one was HCV genotype 2, subgenotype 2c. Most strains carried D310N, an RAS associated with ribavirin, while no RASs associated with sofosbuvir resistance were detected. There is HCV genotype homogeneity with HCV genotype 4, particularly HCV subgenotype 4d, predominating. No major sofosbuvir RASs were found, but ribavirin RAS D310N was common. Future studies should use broader geographic sampling and whole-genome sequencing to comprehensively characterize RASs, their distributions, clinical significance, and temporal trends, thereby guiding treatment strategies in Ethiopia.

**Importance:** Hepatitis C virus infection remains a major global health concern. Its high genetic diversity influences viral evolution, transmission, and response to antiviral therapy. Despite the significant burden of hepatitis C virus infection in sub-Saharan Africa, molecular data on circulating genotypes, subgenotypes, and resistance associated substitutions remain limited. The significance of our research is in characterizing the genetic diversity and resistance associated substitutions among hepatitis C virus strains circulating in Ethiopia. These findings provide important insights into hepatitis C virus molecular epidemiology and may support genomic surveillance, optimization of treatment strategies, and improved understanding of viral evolution trends and patterns among blood donor populations.

## Introduction

Hepatitis C virus (HCV) is one of the major contributors to liver cirrhosis, hepatocellular carcinoma, and viral hepatitis related mortality among hepatotropic viruses (1). An estimated 50 million people are living with the virus globally (2), and Africa bears the third-highest burden of HCV (3). Ethiopia is among the 10 countries that share nearly 2/3 of the estimated global disease burden of viral hepatitis, including HCV (2), and it imposes a significant health and economic burden (4).

Hepatitis C virus (HCV) is a member of the species *Hepacivirus hominis*, genus *Hepacivirus,* family *Flaviviridae*; it is an enveloped, single-stranded, positive-sense RNA (ribonucleic acid) virus. It has ∼ 9.6 kb RNA genome size and a single open reading frame encodes for three structural proteins (Core, Envelop 1, and Envelop 2) and seven non-structural proteins (NS2, NS3, NS4A, NS4B, NS5A, NS5B, and p7) (5).

To date, eight genotypes and 93 subgenotypes of HCV have been identified with distinct geographic variation (6). Additionally, there is a significant difference in genomic structure, pathogenesis, and response to therapy (7). Genotypes 1 and 4 showed limited response to interferon therapy (8), while genotype 3 showed limited susceptibility to direct-acting antiviral agents (DAAs) (9). Moreover, in the clinical context, subtype 1b demonstrated high hepatocellular carcinoma development potential (10), while genotype 3 has a greater tendency for hepatic steatosis (11).

Direct-acting antiviral agents (DAAs) primarily target the non-structural proteins of HCV. Based on their molecular target and mode of action, the existing HCV DAAs are classified into four groups: NS5A inhibitors, NS3/4A protease inhibitors, nucleotide analog inhibitors of NS5B RNA-dependent RNA polymerase (RdRp), and non-nucleoside inhibitors of RdRp (12).

World Health Organization (WHO) recommends Sofosbuvir+ Velpatasvir, Sofosbuvir+ Daclatasvir, and Glecapravir+Pibrentasvir as first-line treatment for HCV (13). This recommendation has been adopted by the Ethiopian treatment guideline, particularly for managing treatment naïve and cirrhosis-free HCV-infected patients. The combination of Sofosbuvir and Ledipasvir is used as an alternative regimen for adults with HCV infection and is preferred for adolescents between the ages of 12-18 years (14).

While DAAs improved HCV treatment, 1– 15% of patients fail to achieve sustained virologic response, resulting from the emergence of resistance associated substitutions (RASs) (15). These RASs were found among 79% to 96% of individuals with first-line treatment and retreatment failure (16). Variants that harbor RASs can alter the shape of a drug target binding site and make it less susceptible to the drug‘s inhibitory activity, subsequently leading to treatment failure (17).

In Ethiopia, 0.93% pooled HCV prevalence was reported among blood donors (18). Blood donors are presumed healthy but may harbor asymptomatic and undiagnosed infections, including HCV, which are capable of transmission through blood transfusion (19). Furthermore, HCV infection in blood donors perhaps reflects the circulating viral strains in the general community; therefore, failure to characterize HCV genetic diversity in this group results in missed opportunities to track circulating genotypes, subgenotypes, and early detection of RASs. However, data regarding the genetic diversity and resistance profile of HCV in Ethiopia is limited, especially among blood donors. The aim of the present study was, therefore, to investigate the genetic diversity of HCV genotypes, subgenotypes, and RASs among blood donors in Addis Ababa using NS5B amplicon based next generation sequencing.

## Materials and Methods

### Study Design, Period, and Setting

Ninety-seven archived serum samples with their complete respective metadata collected from January 2022 to December 2023 from voluntary, apparently healthy blood donors at the Ethiopian Blood and Tissue Bank Service (EBTBS) in Addis Ababa, and who tested positive for HCV antibody by Alinity i chemiluminescent micro particle immunoassay (Abbot diagnostics, Wiesbaden, Germany) tests were included in this study.

### HCV RNA extraction, NS5B gene amplification, and sequencing

HCV RNA was extracted from 200μL of serum using the High Pure Viral nucleic acid kit (Roche v2, Mannheim, Germany). Complementary DNA (cDNA) was synthesized from 10µl of the extracted RNA, random primer at 0.5µM (Invitrogen, Waltham, USA), Superscript IV reverse transcriptase enzyme (Invitrogen, Waltham, USA) at 1.6U/µl, 5× SSIV buffer, ribonuclease inhibitor (Invitrogen, Waltham, USA) at 2U/µl, deoxyribonucleotide triphosphates (dNTPs) at 0.5mM (Invitrogen, Waltham, USA) in final reaction volume of 20µl. The cDNA (5 µl) was added to 15 µL of Platinum SuperFi II Green PCR (polymerase chain reaction) Master Mix (Invitrogen, Waltham, USA) and amplified by nested PCR. The first round PCR with Pr1 (TGGGGATCCCGTATGATACCCGCTGCTTTGA) and Pr2 (GGCGGAATTCCTGGTCATAGCCTCCGTGAA) was performed using the reaction conditions: initial denaturation at 98^0^C for 3 min; 50 cycles of denaturation at 98^0^C for 30 s, annealing at 63^0^C for 30s, and elongation at 72^0^C for 30 s. The final elongation step was at 72^0^C for 10 min (20) and the second round PCR was used by internal primers Pr3 (TATGAYACCCGCTGYTTTGACTC) and Pr5 (GCTAGTCATAGCCTCCGT) under the following condition: 98 ^0^C for 3 min, followed by 35 cycles at 98^0^C for 30 s, 55 ^0^C for 30 s, 72 ^0^C for 30 s, and a final elongation at 72 ^0^C for 10 min (20) and yielded an amplicon size of 381 bp which was visualized by 1% agarose gel. The PCR products were purified using the QIAquick PCR & Gel Cleanup Kit (QIAGEN, Hilden, Germany). Library preparation was performed using the Illumina COVIDSeq RUO Kits (lllumina inc., San Diego, USA) by following the user manual for library preparation. The preparation includes tagmentation of PCR amplicons (fragment and tag PCR amplicons with adaptors), amplify tagmented PCR amplicons (adding dual indexes (i7/i5 and amplifying), pool and clean up, quantify (Quibt ds HS), and normalize library, pool and dilute libraries to 75pM final loading concentration. Sequencing was performed on the Illumina iSeq 100 platform.

### Statistical Analysis

After the data was entered into Excel, data cleaning and validation were done before the analysis. The cleaned data were transported into Stata (v14.2) for analysis. Descriptive results were expressed as percentages for categorical variables.

### Quality Control

Both negative and positive controls were used during extraction, cDNA synthesis, PCR, and library preparation steps. Additionally, the quality and quantity of RNA were checked by nano drop, and all extracts were stored at -80 ^0^C until use.

### HCV genotypes, subgenotypes, phylogenetic, recombination, and RASs analysis

Next-generation sequencing data quality was assessed using FastQC (v0.12.1), and low-quality bases (Phred score < 20 or read length <40 base pairs) and adaptors were trimmed using Fastp (v0.23.4). Dehosting and decontamination were done using Kraken2 (v2.1.3). Post-trimming quality was assessed again using FastQC to check for improvement. High-quality reads were aligned to the reference HCV genome FJ462437 and D50409 using the BWA-MEM (v0.7.17- r1188) algorithm. Samtools (v1.19.2) was used to convert the SAM file into a more compact BAM file format for downstream analysis. Mapping summary metrics, including covered bases, coverage, mean depth, mean base, and mean mapping quality, were summarized or calculated. Finally, consensus sequences of HCV were generated after doing variant calling in BCFtools (v1.21). Then the samples were genotyped by submitting consensus sequences to NCBI BLAST, Geno2Pheno [hcv], and Los Alamos HCV database (https://hcv.lanl.gov/content/index). The genotype results were validated using phylogenetic tree clustering and genetic diversity analyses.

For the phylogenetic analysis, all downloaded HCV strains from the NCBI database, along with strains from the current study (a total of 399 strains) were aligned using the Multiple alignment using Fast Fourier transform (MAFFT) (v7). For tree construction, IQ-TREE was used. A phylogenetic tree was then constructed using the Maximum Likelihood method. The robustness of the tree topology was assessed with 1000 bootstrap replicates. Finally, the resulting phylogenetic tree in Newick format was imported into the Interactive Tree of Life (iTOL) platform for annotation and visualization.

All strains that were used for the phylogenetic tree construction were reused again for recombination analysis using RDP5 (Recombination Detection Program). Recombination events detected by at least four independent methods (RDP, GENECONV, BootScan, MaxChi, Chimaera, SiScan, and 3Seq/Topol) were considered. All analyses were performed using the default parameters in RDP5. The RASs analysis for sofosbuvir, and ribavirin web-based tool Geno2Pheno [hcv] was used.

### Data availability

The HCV NS5B partial gene sequences generated in this study are publicly available in the NCBI GenBank database under accession numbers PZ540809-PZ540843. The raw sequencing reads supporting these findings have been deposited in the NCBI Sequence Read Archive (SRA) under (SRR39009979, SRR39009978, SRR39009976, SRR39009975, SRR39009974, SRR39009973, SRR39009972, SRR39009971, SRR39009970, SRR39009969, SRR39009968, SRR39009967, SRR39009965, SRR39009964, SRR39009963, SRR39009962, SRR39009961, SRR39009960, SRR39009959, SRR39009958, SRR39009957, SRR39009956, SRR39010065, SRR39010064, SRR39010063, SRR39010062, SRR39010061, SRR39010060, SRR39010059, SRR39010058, SRR39010057, SRR39010056, SRR39010054, SRR39010053, and SRR39010052), linked to BioProject PRJNA1475203.

## Results

### Demographic and donor characteristics of anti-HCV positive study participants

Among the 97 anti-HCV positive samples, one sample showed co-infection with syphilis. The age of blood donors ranged from 19 to 60 years, with a median of 27 years (Interquartile range: 22- 37). Male donors represented 56 (57.7%). Regarding blood group distribution, “O” was most common (31.96%), followed by “A” and “B” (29.90% each). More than half of the donors (50.52%) had previously donated blood, and the majority were Rh-positive (91.75%). A total of 45 samples were PCR-positive, yielding a PCR positivity rate of 46.39% (**Table 1**).

**Table 1:** Demographic and donor characteristics of the anti-HCV positive blood donors, Addis Ababa, Ethiopia, from January 2022 to December 2023 (n = 97).

| Variable | Category | Total number (%) | PCR result |  |
| --- | --- | --- | --- | --- |
|  |  |  | Positive | Negative |
| Gender | Male | 56(57.73) | 28 | 28 |
|  | Female | 41(42.27) | 17 | 24 |
| Age | 19-29 | 57 | 21 | 36 |
|  | 30-39 | 18 | 8 | 10 |
|  | ≥40 | 22 | 16 | 6 |
| Blood group | A | 29(29.9) | 16 | 13 |
|  | B | 29(29.9) | 11 | 18 |
|  | AB | 8(8.24) | 4 | 4 |
|  | O | 31(31.96) | 14 | 17 |
| Rh factor | Positive | 89(91.75) | 41 | 4 |
|  | Negative | 8(8.25) | 4 | 4 |
| Donation history | First time donor | 48(49.48) | 18 | 30 |
|  | Repeated donor | 49(50.52) | 27 | 22 |
| RDT result | Positive | 11(11.34%) | 9 | 2 |
|  | Negative | 86(88.66) | 36 | 50 |
| **Abbreviation:** RDT = Rapid diagnostic test for HCV (One-step test device HCV, EGENS Diagnostics) |  |  |  |  |
| **Note:** Each variable proportion among PCR positive and negative was determined using cross-tab in Stata Statistical Software (version 14). |  |  |  |  |

### Sequencing and NGS analysis of HCV strains

Of the 45 PCR-positive samples, 40 were successfully purified and sequenced. Sequencing success rate was 87.5% (35/40). Five sampl2es were excluded due to low quality, insufficient coverage, or inadequate sequencing depth (< 30x). A total of 35 high-quality samples were obtained with a fragment length ranging from 40 to 150 nucleotides. The mean mapped region coverage and mean depth of coverage was 79.54% and 5938.04, respectively.

### HCV genotypes, subgenotypes, phylogenetic, and recombination analysis

To characterize the circulating HCV genetic diversity and identify RASs, the NS5B region of HCV was sequenced from antibody-positive blood donor samples. Both online databases and Phylogenetic analysis were performed to determine genotypes and subgenotype distributions. Genotypic analysis of 35 HCV sequences from blood donors identified two genotypes: genotype 4 was predominant in 34 (97.14%) samples, while genotype 2 was detected in 1 sample (2.86%). Subgenotype analysis revealed that all genotype 4 strains were 4d (100%, n = 34), whereas the single genotype 2 strain was 2c.

Phylogenetic analysis further confirmed HCV typing tools results, with strains from this study clustering with the previously described HCV genotype 4 and 2. All HCV genotype 4 strains (34 strains) from the current study formed a distinct cluster with previously identified subgenotype 4d strains, showing the highest similarity to strains from Ethiopia and the United States of America (USA). The single genotype 2c strain clustered within a separate, well-supported clade alongside genotype 2c strains from Belgium and Denmark (Figure 2). Moreover, all HCV strains from this study underwent recombination analysis using RDP5, but none showed evidence of recombination.

**Figure 1:**
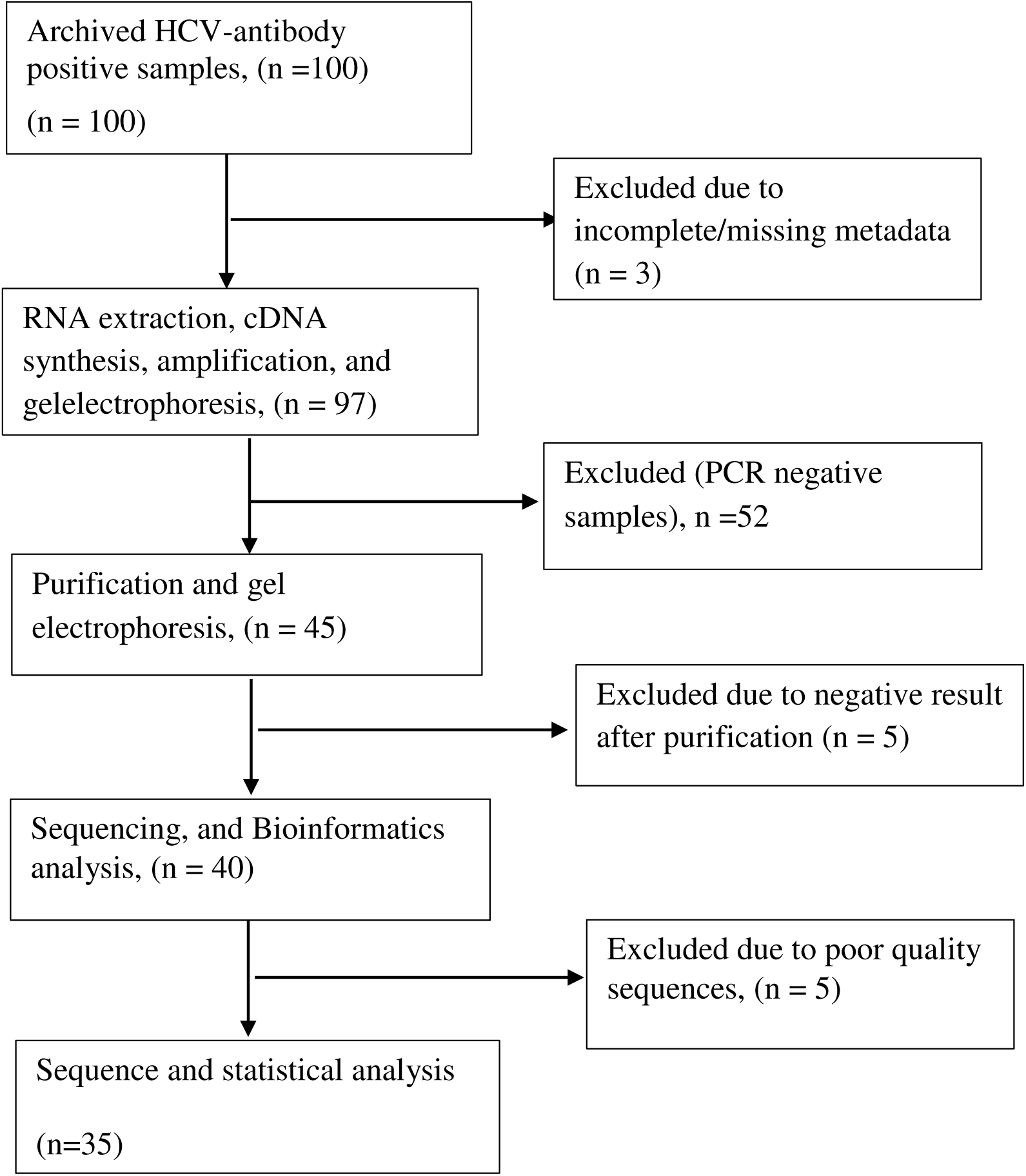
Workflow of HCV from data cleaning to result interpretations of anti-HCV positive blood donors, Addis Ababa, Ethiopia, from January 2022 to December 2023 (n = 97) Of 100 archived HCV antibody-positive samples, conventional PCR was done for 97 samples, which have complete metadata. The PCR result yielded 45 positives, and the remaining 52 PCR-negative samples were removed. Of the 45 PCR-positive samples, 5 were removed, as they became negative after purification. A total of 40 samples were sequenced, resulting in 35 consensus sequences for genotyping and RASs analysis.

**Figure 2:**
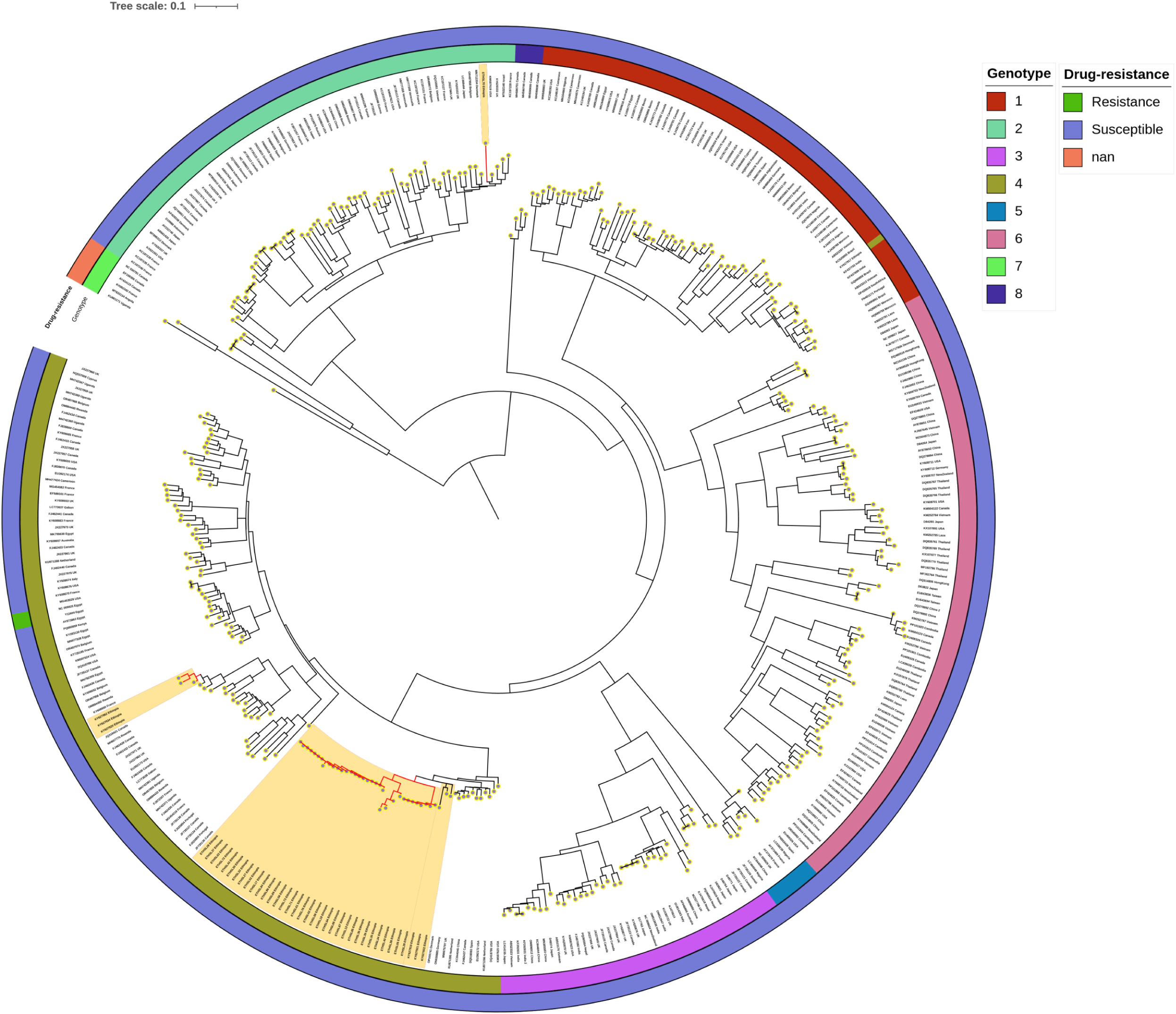
Phylogenetic tree of HCV strains from anti-HCV positive blood donors, Addis Ababa, Ethiopia, from January 2022 to December 2023 (n = 35). A Phylogenetic tree was constructed based on the partial HCV NS5B gene, using 35 strains from the present study along with 364 GenBank sequences. The ETHSL represents samples from the present study, then the sample identification number, and the country of origin (e.g., ETHSL13 Ethiopia). Strains from the GenBank database are designated by their accession number and their respective country of origin (e.g, KU861171 Uganda). Bootstrap values ≥ 90 are indicated at the nodes. Branch lengths are proportional to sequence divergence. Similar geometric shapes within a node denote that they belong to one family. The two rings represent the drug resistance profile and genotype, respectively, from outer to inner. Strains highlighted with a red branch and shaded with a light yellow color represent strains from Ethiopia, including strains from the present study. The phylogenetic tree analysis revealed that all genotype 4 strains are subgenotype 4d, and a single genotype 2 strain that belongs to subgenotype 2c. Ethiopian strains (ETHSL) from the current study formed a highly supported monophyletic cluster, indicating close evolutionary relatedness and suggesting circulation of genetically conserved viral variants within the Addis Ababa blood donors. The short internal branch lengths observed among the majority of the Ethiopian strains show low genetic divergence and support the presence of limited sequence differences among these strains. The highly genetically related or monophyletic nature of the phylogenetic tree was further confirmed by calculating the overall mean distance between subgenotype 4d strains from the present study: mean ± standard error (5% ± 1%).

### RASs in the NS5B gene of HCV Strains

To identify RASs in the partial NSB gene an online database i.e Geno2Pheno [hcv] was used. All 35 HCV strains harbored multiple RASs in the NS5B gene, ranging from a minimum of three to a maximum of seven per strain, and among observed strains, 18 substitutions were unique. Although no well-known sofosbuvir RASs (S282T, L320F/C, C316Y/N) were detected, several NS5B mutations with no prior association to resistance were observed. In addition, the RAS D310N was detected in the majority of the strains (22/35, 62.86%) (Table 2).

**Table 2:**
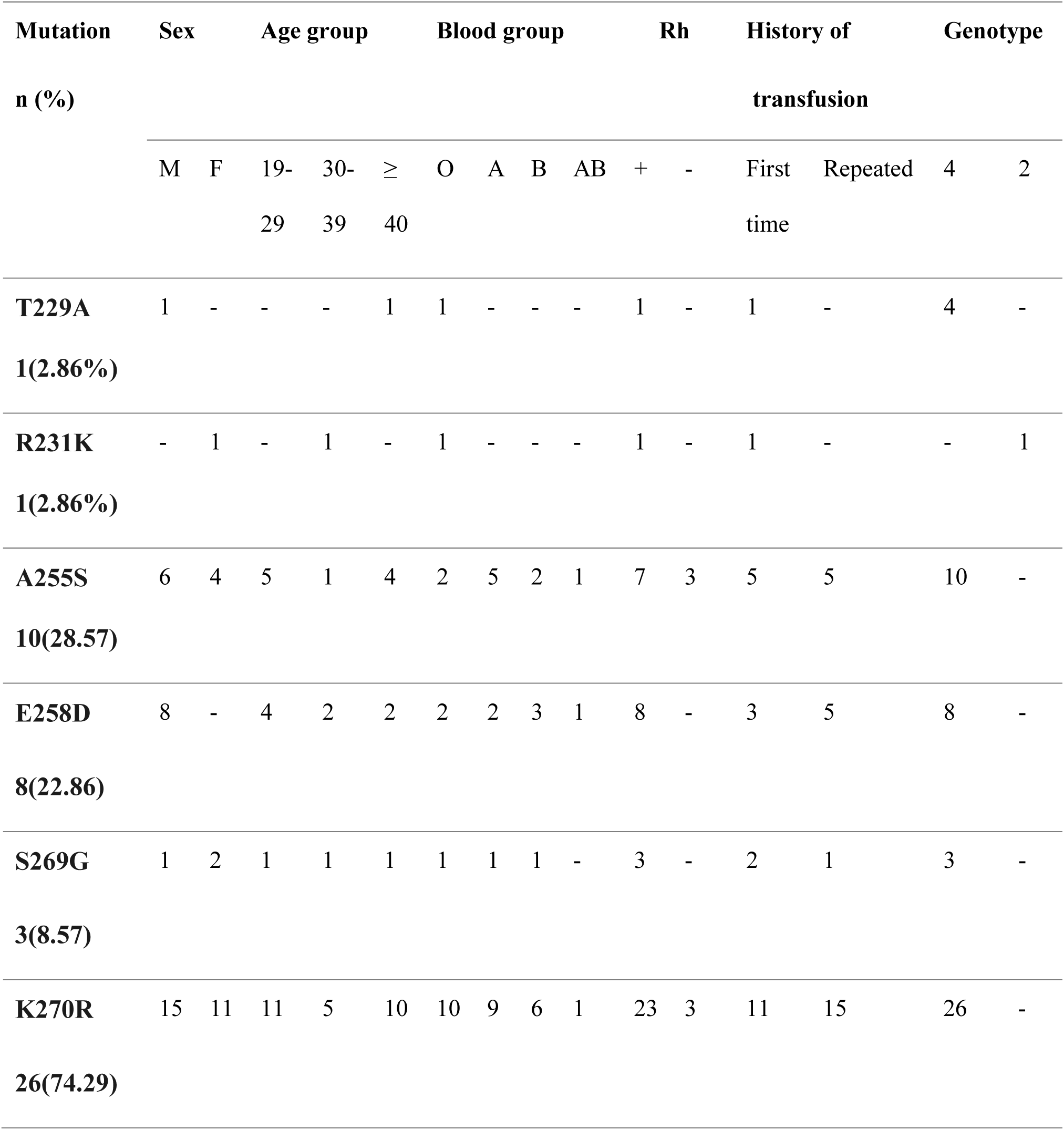
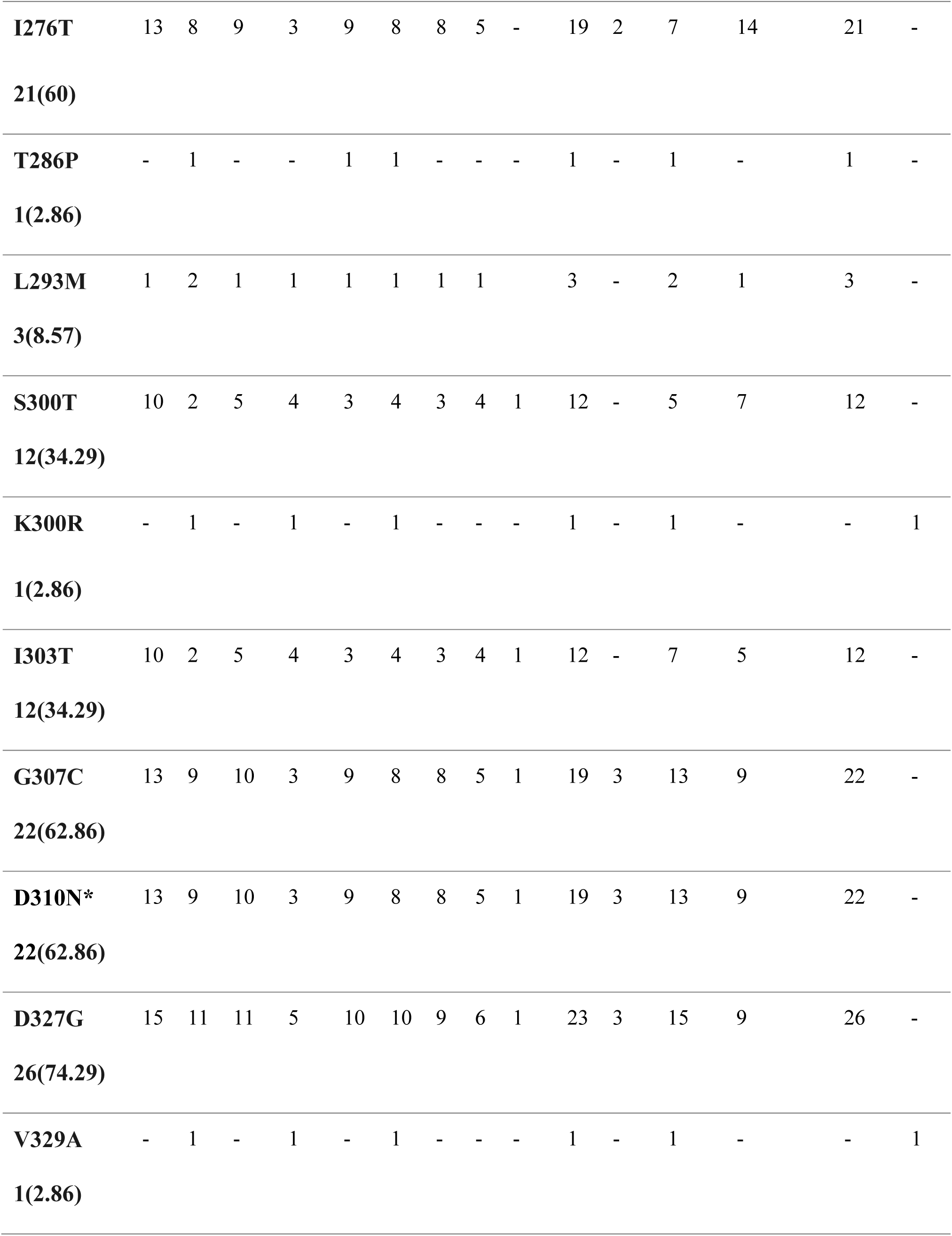
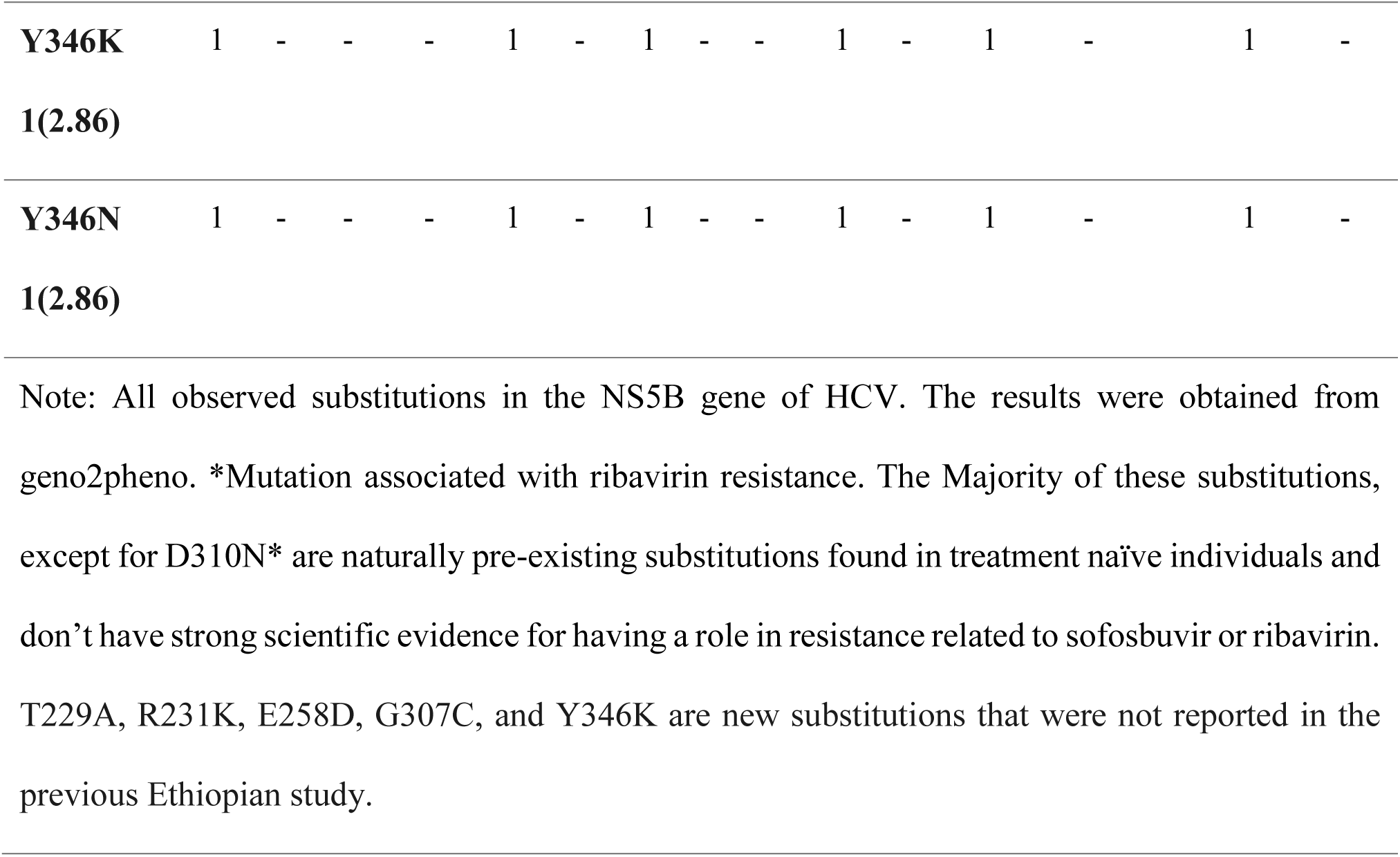
List of mutations identified in the partial NS5B gene of anti-HCV positive blood donors, Addis Ababa, Ethiopia, from January 2022 to December 2023 (n = 35).

## Discussion

In our analysis, we identified predominant HCV genotype 4 strains (n = 34, 97.14%), followed by a single genotype 2 strain. Previously reported less common genotypes 1, 3, and 5 were not detected. The predominance of genotype 4 in the current study is concordant with earlier Ethiopian studies, which reported 77.6% genotype 4 prevalence among blood donors (21). Moreover, predominantly circulating genotype 4 was not only identified in the blood donor population but also in different populations (chronic patients, HIV-HCV co-infected, and HCV mono-infected), and across different time periods (22–24). In agreement with our finding, a study from the Democratic Republic of Congo blood donors found genotype 4 (50%) as a dominant strain (25). In contrast to our result, predominant genotype 1 (77.5%) was found in Brazilian (26), 58.1% in the USA (27), 58.68% in China (28), and 69.05% in Indian (29) blood donors. Different genotype distribution patterns across geographical areas are a possible reason for the observed difference (30).

HCV genotype 4 is believed to have originated in Central Africa and subsequently spread to North Africa and other continents due to human migration and major population-changing events such as World War II (31). The predominance of genotype 4 in Ethiopian studies suggests that this genotype has been endemic for a longer time in the Ethiopian population (21). From a clinical perspective, this genotype is among the genotypes associated with insulin resistance (32). The proposed mechanism for the observed insulin resistance in HCV can be explained by the HCV core protein, which increases serine phosphorylation of insulin receptor substrate through downregulation of the Tuberous Sclerosis Complex 1 or 2 (TSC1/TSC2) complex, with subsequent upregulation of the mammalian target of rapamycin or ribosomal protein S6 kinase 1 (mTOR/S6K1) (33).

The subgenotype analysis revealed that all genotype 4 strains were subgenotype 4d, and the genotype 2 strain was subgenotype 2c. Although the dominance of subgenotype 4d is supported by two studies in Ethiopia (21, 22), the divergence of subgenotype 4 was low compared to the previous study (21). The observed differences are likely to be attributed to variations in study populations, as the previous study included multiple geographically distinct regions (Gondar, Mekelle, Adama, and Jimma), potentially capturing greater genetic diversity.

In the current study, the prevalence of subgenotype 4d increased markedly from the previous report, rising from 34.7% (21) to 97.14%. Notably, nearly a decade has elapsed since the earlier study, highlighting temporal changes in subgenotype distribution. Phylogenetic analysis revealed that, except for a single strain, all genotype 4d sequences clustered closely with previously identified Ethiopian 4d strains. This near-exclusive 4d predominance may reflect a founder effect and localized transmission dynamics within the study population, suggesting that a specific subgenotype has persisted and expanded over the past ten years.

Absence of RASs for DAA drug sofosbuvir in the present investigation is supported by local and international studies. A study from Ethiopia reported no RAS to sofosbuvir (21). Moreover, studies consistently reported very low prevalence of RASs to sofosbuvir across different genotypes and different study populations (34, 35). High genetic barrier, and classic mutation S282T resulted in replication-incompetent, which are biologically plausible reasons for the low detection of RASs (36).

The Ethiopian viral hepatitis treatment guideline recommends ribavirin in combination with other DAAs in different populations, including cirrhotic patients (14). In the present study, 22 (62.86%) of the analyzed strains harbored RAS to ribavirin. This proportion is higher than that reported in previous studies; an earlier Ethiopian study detected ribavirin-related RASs in only 2 of 46 strains (4.35%) (21). Similarly, low prevalence was reported among treatment naïve patients from Cameroon (79, 8.54%) (37), Vietnam (2/100, 2%) (38), and in Brazil (5, 7.2%) (39). A Possible explanation for the observed high RAS in the present finding might be due to differences in genotype distributions. Unlike studies with a low prevalence of RAS for ribavirin, our study is dominated by genotype 4, which might explain the observed variation (40). Likewise, the D310N substitution showed differential distribution across subgenotypes (1a, 1b, 2a, 2b, and 3a), with subgenotype 3a accounting for the highest proportion (92.9%) in a study conducted in Canada (41).

### Study Strengths and Limitations

This study has two key strengths. First, to the best of our knowledge, it is the first study of HCV in Ethiopia conducted entirely within a national laboratory setting, from sample processing to molecular analysis. Second, the study employed NGS, which offers higher sensitivity, greater accuracy, and comprehensive genomic coverage compared with Sanger sequencing.

This study has limitations that should be acknowledged. First, sequencing was a partial region of the NS5B gene, which precluded a more comprehensive assessment of viral diversity and RASs. Second, the absence of viral load determination limited our ability to correlate genotypes and clinically relevant mutations with viral replication levels, which could have provided further clinical and epidemiological insights.

### Conclusions

There is HCV genotype homogeneity with predominance of HCV genotype 4 and subgenotype 4d among blood donors in Addis Ababa. A notable increase in HCV genotype 4 was observed compared to previous study. Overall, the RASs to sofosbuvir were negligible. In contrast, a high prevalence of RAS for ribavirin was observed among HCV antibody-positive blood donors in Addis Ababa. Further, a large-scale study is warranted to better understand the profile and impact of genetic mutations, including RASs, in HCV.

## Supporting information

**Table S1.** Metadata of HCV NS5B sequences used for phylogenetic analysis, including accession numbers, genotypes, subgenotypes, and resistance profiles.

**Table S2.** Summary of Coverage, depth, and quality metrics of HCV NS5B sequencing data from the present study

## Acknowledgment

The authors would like to acknowledge the Armauer Hansen Research Institute (AHRI) for financial support for sample collection and laboratory reagents. The authors also extend their gratitude to the University of Gondar (UoG) for sponsorship and a monthly stipend for the student (Helen Terefe). Finally, our appreciation goes to the EBTBS and the study participants.

## Declarations

### Author Contributions

**Conceptualization:** Helen Terefe, Andargachew Mulu, Adane Miheret, and Dawit Hailu Alemayehu. **Data curation**: Helen Terefe, Gizachew Gemechu, Dawit Hailu Alemayehu**. Formal analysis:** Helen Terefe, Abaysew Ayele**. Investigation:** Helen Terefe**. Methodology:** Helen Terefe, Dawit Hailu Alemayehu, Andargachew Mulu, Tesefaye Gelanew, Adane Miheret, and Yimer Demisse**. Project administration:** Andargachew Mulu, Dawit Hailu Alemayehu, and Helen Terefe. **Resources:** Dawit Hailu Alemayehu, Abaysew Ayele, and Helen Terefe. **Review and editing:** Andargachew Mulu, Tadelo Wondemagege, Abaysew Ayele, Adane Miheret, Tesefaye Gelanew, Dawit Hailu Alemayehu, Gizachew Gemechu, and Gashaw Adane. **Software:** Abaysew Ayele and Helen Terefe **Supervision:** Andargachew Mulu, Tesefaye Gelanew, Adane Miheret, Tadelo Wondemagege, Gashaw Adane. **Validation:** Helen Terefe, Dawit Hailu Alemayehu, Abaysew Ayele**. Visualization:** Abaysew Ayele and Helen Terefe. **Writing original draft preparation:** Helen Terefe

### Ethical consideration

The study received ethical approval from both the School of Biomedical and Laboratory Sciences (SBLS), College of Medicine and Health Science, University of Gondar (SBLS/4477/), and AHRI/All-Africa Leprosy Tuberculosis and Rehabilitation Training Center(ALERT) Ethical Review Committee (Protocol number: (PO-109-25)). A permission letter was obtained from AHRI Bio Bank so as to access samples. Blood donors have already given their informed consent as per the EBTBS standard donor questionnaire for further research works; consequently, we were granted a waiver of informed consent for this research.

## Abbreviations and acronyms

The following abbreviations and acronyms are used in this manuscript:

AHRI: Armauer Hansen Research Institute
cDNA: Complementary DNA
dNTP: Deoxyribonucleotide triphosphates
DAA: Direct-acting antiviral agent
EBTBS: Ethiopian Blood and Tissue Bank Service
HCV: Hepatitis C virus
MAFFT: Multiple alignment using Fast Fourier transform
NS5B: Non-structural protein 5B
PCR: Polymerase Chain Reaction
RAS: Resistance-associated substitution
RNA: Ribonucleic acid
USA: United States of America

